# Host fruit as a suitable bacteria growth substrate that promotes larval development of *Bactrocera dorsalis* (Diptera: Tephritidae)

**DOI:** 10.1101/834119

**Authors:** Mazarin Akami, Xueming Ren, Yaohui Wang, Abdelaziz Mansour, Shuai Cao, Xuewei Qi, Albert Ngakou, Chang-Ying Niu

**Author notes:** Authors for correspondence: CYN.

## Abstract

The ability of a host plant to act as a substrate or media for larval development may depend on how good it is at offering suitable nutrients for bacterial growth. In this study, we hypothesized that the suitability of a fruit type for fruit fly larval development is positively correlated with the ability of that fruit to act as a substrate/media for fruit fly symbiotic bacterial growth. We allowed a single female fruit fly to lay eggs on five different host fruits, then we monitored the larval development parameters across five generations and analyzed the bacterial community structure of larvae developing in 2 of these hosts (apple and banana) at the first and fifth generations. Results indicate that the larval length and dry weight did not vary significantly across experimental generations, but were greatly affected by fruit types and larval stages. The larval development time was extended considerably in apple and tomato but shortened in banana and mango. There was a significant shift in bacterial community structure and composition across fruits and generations. The bacterial community of larvae within the same fruit (apple and banana) clustered and was similar to the parental female (with the predominance of Proteobacteria), but there was a shift at the fifth generation (dominance of Firmicutes). Banana offered a suitable better development and growth to larvae and bacteria, respectively, compared to apple in which reduced larval development and bacterial growth were recorded. Although additional experiments are needed to adequately show that the differences in microbiome seen in fruit fly larval guts are the actual driver of different developmental outcomes of larvae on the different fruits, at the very least, our study has provided intriguing data suggesting interaction between the diets and gut microbial communities on insect development.

**Importance and Significance of the study:** Tephritid fruit flies entertain complex interactions with gut bacteria. These bacteria are known to provide nutritional benefits to their hosts, by supplementing missing nutrients from the host diets and regulating energy balance. Foraging for food is a risky exercise for the insect which is exposed to ecological adversities, including predators. Therefore, making beneficial choice among available food substrates is a question of survival for the flies and bacteria as well. Our study demonstrates interactions between the host fly and its intestinal bacteria in sustaining the larval development while foraging optimally on different fruit types. These findings add a novel step into our understanding of the interactions between the gut microbial communities and *B. dorsalis* and provide avenues for developing control strategies to limit the devastative incidence of the fly.

## Introduction

Gut bacteria and fruit flies are thought to have a close evolutionary and biological relationships with a wide degree of interdependence [1–3]. This interaction shapes the host fitness and the abundance of gut microbiota [2], and is modulated by the availability and nutritional quality of host fruits. Different fruit differ in their suitability for fruit fly larvae. If fruit fly larvae gain many of their primary nutritional needs from bacterial break-down products, then maybe the quality of fruit for fruit fly larvae depends on how good the host is for bacterial growth and survival, with the fruit’s value to fruit fly larvae being of secondary importance.

Some tropical fruits such as banana and apple have been shown to contain a good amount of pectin intensively used as food additive [4–6]. Pectin from fruit has been reported to be involved in defense mechanisms against external aggressions and plant pathogens [7]. In addition, fruits are generally low in protein content. The larvae of the oriental fruit fly *Bactrocera dorsalis* require a large amount of sugar and nitrogen content diet for their growth [8]. In order to develop in nutritionally poor fruits, gut bacteria may come into play in supplying missing sugar metabolites and marginal amino acid residues from the various fruit types to sustain the larval development [9,10].

The ability of gut bacteria to modify the diet composition [11] and host transcriptome [12] allows aphids to survive on plants with poor nutrient values [13], allows the higher termites *Nasutitermes takasagoensis* to digest cellulose from wood [14], *Bombyx mori* to degrade pectin from mulberry leaves [15] and *Anoplophora glabripennis* to feed on multiple hosts by disrupting the expression levels of numerous genes involved in digestion and detoxification [12]. Insects are exposed to several ecological adversities (predators and abiotic factors) and the gut bacterial isolates were conjected to help *B. dorsalis* to making beneficial compromises between the feeding time, nutrient ingestion and fitness [16,17]. These ecological compromises (nutritional tradeoffs) result in differences in *B. dorsalis* gut microbiome, whose community structure and diversity vary or shift from one fruit to the other in response to the nutritional adaptation, not only for the host but also for the gut microbiotas. In this event, *B. dorsalis* may rely on its gut-associated bacteria for fitness and survival, which in turn may shape the foraging of the host according to their nutritional requirements [16,17].

In this study, we evaluated the extent of the interactions between the diets and the microbial communities on larval development. We predicted that the suitability of a fruit type for fruit fly larval development is positively correlated with the ability of that fruit to act as a substrate/media for fruit fly symbiotic bacterial growth. The method consisted of allowing a single female fly to lay eggs on five different fruit types for 24h and monitoring the larval population dynamics across five generations. At each generation, the larval development parameters (length, weight and development time) were evaluated within and between fruit types. We thus presumed that larval development parameters would be highly enhanced in fruits, thus offering optimal nutrients for development [18]. Data generated from this study allowed us to conject that the gut microbiome can shift its population in response to nutritional adaptations of fly larvae.

## Materials and methods

### Insect rearing and maintenance

All experiments were conducted in a controlled environment (25±1.5°C, 65±10% RH and 16:8 light: dark cycle). Newly emerged lab-reared flies were fed artificial full diet consisting of Tryptone (25 g/L), Yeast extract (90 g/L), Sucrose (120 g/L), Agar powder (7.5 g/L), Methyl-p-hydroxybenzoate (4 g/L), Cholesterol (2.3 g/L), Choline chloride (1.8 g/L), Ascorbic acid (5.5 g/L) in 1 L of distilled water.

### Host fruits preparation

Ripe fruits (mango, banana, apple, citrus, and tomato) were bought from a local supermarket (Wuhan, China) and surface sterilized by soaking them in 2% sodium hypochlorite for 20 minutes and rinsed with deionized distilled water. The sterilized fruits were air dried under a laminar flow hood to avoid airborne contamination. Five of each type of fruit were needed for the bioassays (therefore the total number of fruit were 5 × 5 = 25), and five replicates were used for each fruit type (125 fruits in total).

### Bioassays

One mating couple (11-15 day-old) was extracted from the rearing cage and introduced into a new cage (15×15×15cm). The different fruits were separately added into the cage in which the single female was allowed to lay eggs for 24 hours before replacing the fruit with another one. The process was repeated for the five fruit types (Fig 1).

**Figure 1.**
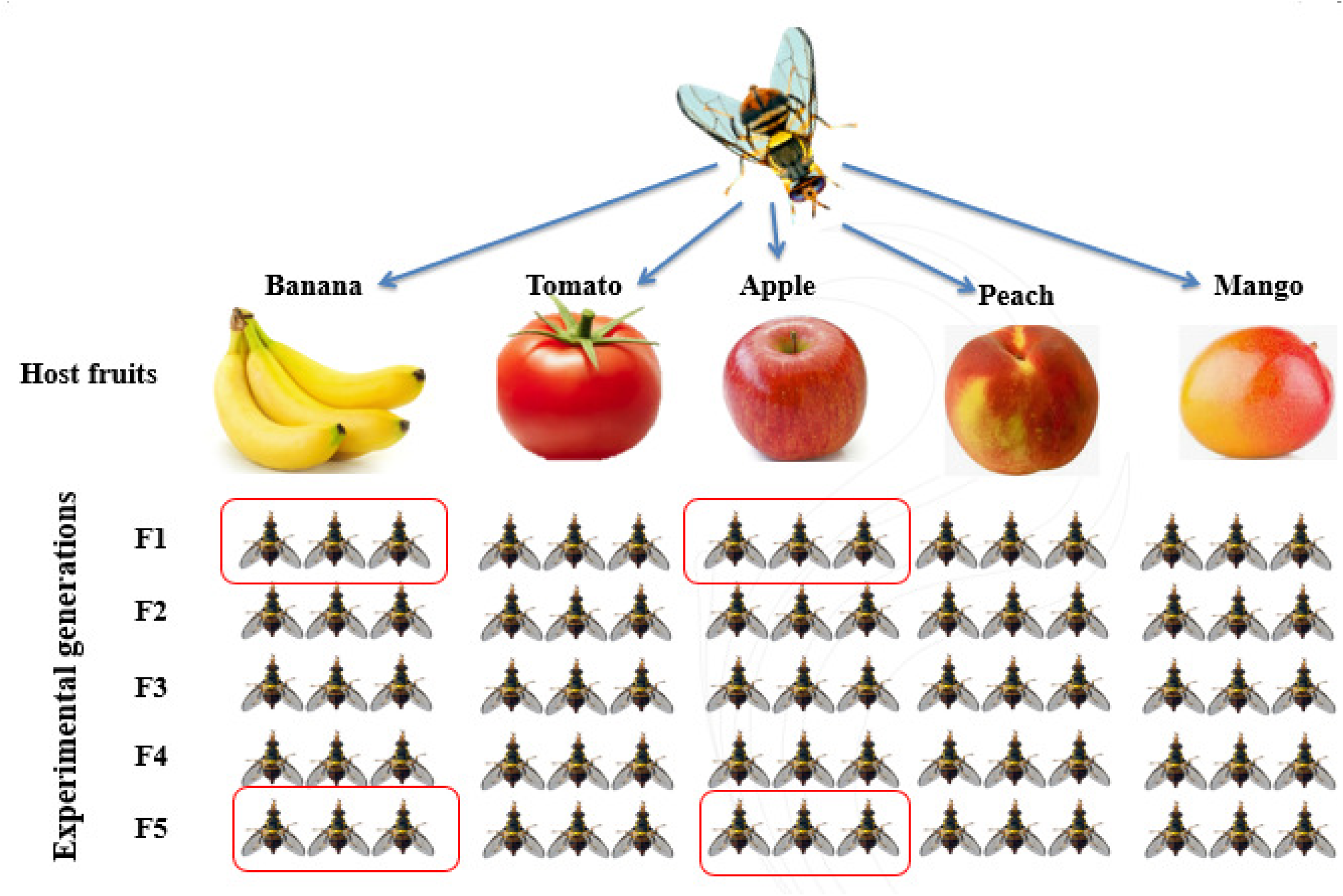
Experimental design. The highlighted ones are samples used for microbial analyses.

To determine how bacteria and fruit type affect larval development, the larval developmental parameters were monitored in all fruit types following the oviposition, starting from the parental female. Fruits bearing eggs were incubated individually in 1L transparent plastic cups and sealed with a fine mesh (25±1.5°C, 65±10% RH and 16:8 light: dark cycle). Eggs and developing larvae were extracted periodically from each fruit type at 3, 6 and 9 days (corresponding to the 1st, 2nd, and 3rd instar larvae). The body length of the extracted larvae (to the nearest 0.03 mm), the larval dry weight and developmental time were measured before the larvae were anesthetized in cold 95% ethanol. All Larvae (15) were kept in 95% ethanol and were either used immediately or preserved at −80°C until used for further analyses (measurement of larval length, dry weight and microbiome analyses). The remaining larvae from each fruit were allowed to develop till adult emergence. The adults that emerged were maintained under artificial diet as described above till the flies reached sexual maturity. A single female from the cohort of each fruit was allowed to lay eggs on new fruit to produce subsequent generation and the process was repeated till five generations.

### Microbial analyses

#### Genomic DNA extraction and amplicon generation

Total genome DNA from samples was extracted using the CTAB/SDS method [19]. DNA quality was checked on 1% agarose gels using a ladder, and the purity was checked as above. DNA was diluted to 1ng/µL with sterile distilled water.

The V1-V3 variable region of the bacterial 16S rDNA gene was amplified to construct a gene library using bar-coded and broadly conserved primers for the PCR reaction: 27F_5′ <u>CCTATCCCCTGTGTGCCTTGGCAGTCTCAG</u>AGAGTTTGATCCTGGCTCAG-3′, and 533R_5′-<u>CCATCTCATCCCTGCGTGTCTCCGACGACT</u>NNNNNNNNTTACCGCGGCTGCTGCAC-3′ [20]. This contains the A and B sequencing adaptors (454 Life Sciences) to facilitate pooling, segregation, sequencing and amplification of ~536 bp region of the mentioned gene (“Ns” represent the 8 nt barcode sequence for multiple samples while the underlined sequences represent the A-adaptor). All PCR reactions were carried out with Phusion® High-Fidelity PCR Master Mix with GC Buffer (New England Biolabs, Ipswich, MA, USA) and high-fidelity polymerase (New England Biolabs).

The PCR conditions were as follows: initial denaturation at 94°C for 2 min, followed by 30 cycles of denaturation at 94°C for 1 min, annealing at 60°C for 30 s, and extension at 72°C for 1 min and a final extension step of 10 min at 72°C. PCRs of DNA-free samples were run to check potential contamination of buffers and primers [21]. PCR amplicons were later subjected to electrophoresis on a 2% agarose gel, stained with ethidium bromide, and the targeted fragment size (400-450bp) was extracted, purified with Qiagen Gel Extraction Kit (Qiagen, Germany) and quality checked before pyrosequencing. Amplicon libraries were generated using TruSeq® DNA PCR-Free Sample Preparation Kit (Illumina, USA) following the manufacturer’s recommendations and index codes were added. Library quality was assessed on the Qubit@ 2.0 Fluorometer (Thermo Scientific) and Agilent Bioanalyzer 2100 system using a DNA1000 lab chip (Agilent), respectively. The library was then amplified by emulsion PCR before 454 pyrosequencing was performed from the A-end on an Illumina HiSeq2500 platform using a GS FLX Titanium system according to the manufacturer’s instructions (Roche 454 Life Sciences) and 250 bp paired-end reads were generated.

#### Bioinformatics analyses

Paired-end reads were assigned to samples based on their unique barcode and truncated by cutting off the barcode and primer sequence. Paired-end reads were merged using FLASH 1.2.7 [22]. Quality filtering on the raw tags was performed under specific filtering conditions to obtain the high-quality clean tags [23] using QIIME1.7.0 [24] quality controlled process (Edgar et al. 2011). The tags were compared with the reference database using UCHIME algorithm to detect and remove chimera sequences before obtaining active tags [25].

Operational taxonomic units (OTUs) analyses were performed by UPARSE 7.0.1001 [26]. Sequences with ≥97% similarity were assigned to the same OTUs. The representative sequence for each OTU was screened for further annotation, and the GreenGene Database [27] was used based on the Ribosomal Database Project (RDP) classifier 2.2 algorithm [28] to annotate the taxonomic information. OTUs abundance information was normalized using a standard of sequence number corresponding to the sample with the least sequences. The Good’s coverage, the abundance-based coverage estimator (ACE), the bias-corrected Chao1 richness estimator, the jackknife estimator of species richness, and the Shannon-Weaver and Simpson diversity indices, were calculated with the Mothur package.

#### Statistical analyses

All data on developmental parameters (larval length, larval dry weight and development time) were tested for homogeneity of variances using Levene’s tests. The crucial factors that affect the developmental parameters were checked by using the regression model (SPSS) with host fruits and experimental generations as effects. The one-way analysis of variance (ANOVA) was used to analyze differences in developmental data and sequencing data. New Duncan’s Multiple Range Test (NDMRT) test at P = 0.05 significance, was used for mean separations within and between samples. The results of sequencing parameters were presented as means of the three biological replicates. Statistical analyses were carried out using SPSS 20.0 software (Statsoft Inc, Carey, J, USA). OriginPro 8.5.1 software was used to construct graphs. We used the Analysis of similarity (ANOSIM) to determine whether the differences of *B. dorsalis* bacterial community between treatments (P, A1, A5, B1 & B5) are significantly higher than those in the group and whether the grouping is meaningful.

## Results

### Larval development parameters

#### Larval length

The larval length and dry weight were not affected by the experimental generations (Regression Model, F = 146.194; df = 4; R^2^ = 0.857; t = 5.013; P = 0.236 and F = 257.234; df = 4; R^2^ = 0. 798; t = 7.835; P = 0.184, respectively, but were significantly affected by fruit types and larval stages (Regression Model, F = 71.217; df = 4; R^2^ = 0.958; t = 1.464; P ˂ 0.0001 and F = 71.217; df = 2; R^2^ = 0.958; t = 1.464; P ˂ 0.0001, respectively) (Fig 2 & 3).

**Figure 2.**
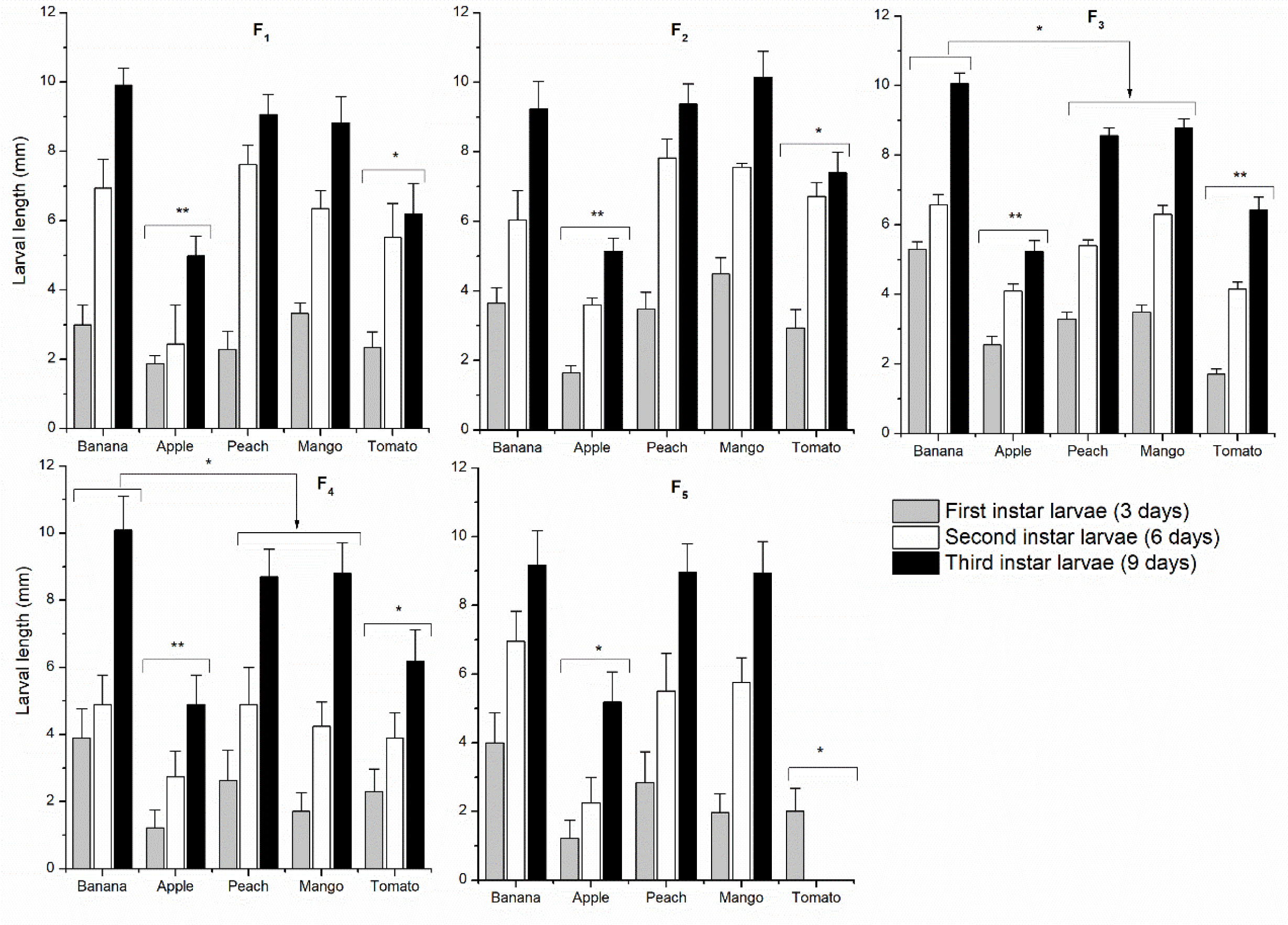
Effect of host fruits suitability on *B. dorsalis* larval growth across five generations.

The larval growth increased significantly across stages in all the experimental fruits and generations (Regression Model, F = 116.643; df = 2, 4; R^2^ = 0.837; t = 7.322; P ˂ 0.0001) and the highest larval length recorded in Banana, peach and mango in comparison to apple and tomato (ANOVA, F = 70.007; df = 4; P ˂ 0.0001 and F = 139.185; df = 4; P ˂ 0.001, respectively) (Fig 2). Except for tomato whose larval development stopped at the first instar of the fifth generation, all the host fruits allowed complete larval development up to the fifth generation (Fig 2).

#### Larval weight

The larval growth positively correlated with larval dry weight across generations in all host fruits (R^2^ = 0. 978; P ˂ 0.0001) (Fig 3) and the two parameters were proportional to each other. Moreover, a paired analysis of larval growth and dry weight revealed a significant interaction between the two parameters in selecting the suitability of host fruits by *B. dorsalis* (F = 40.074; df = 1, 4; R^2^ = 0.9984; P ˂ 0.0001) (Fig 3).

**Figure 3.**
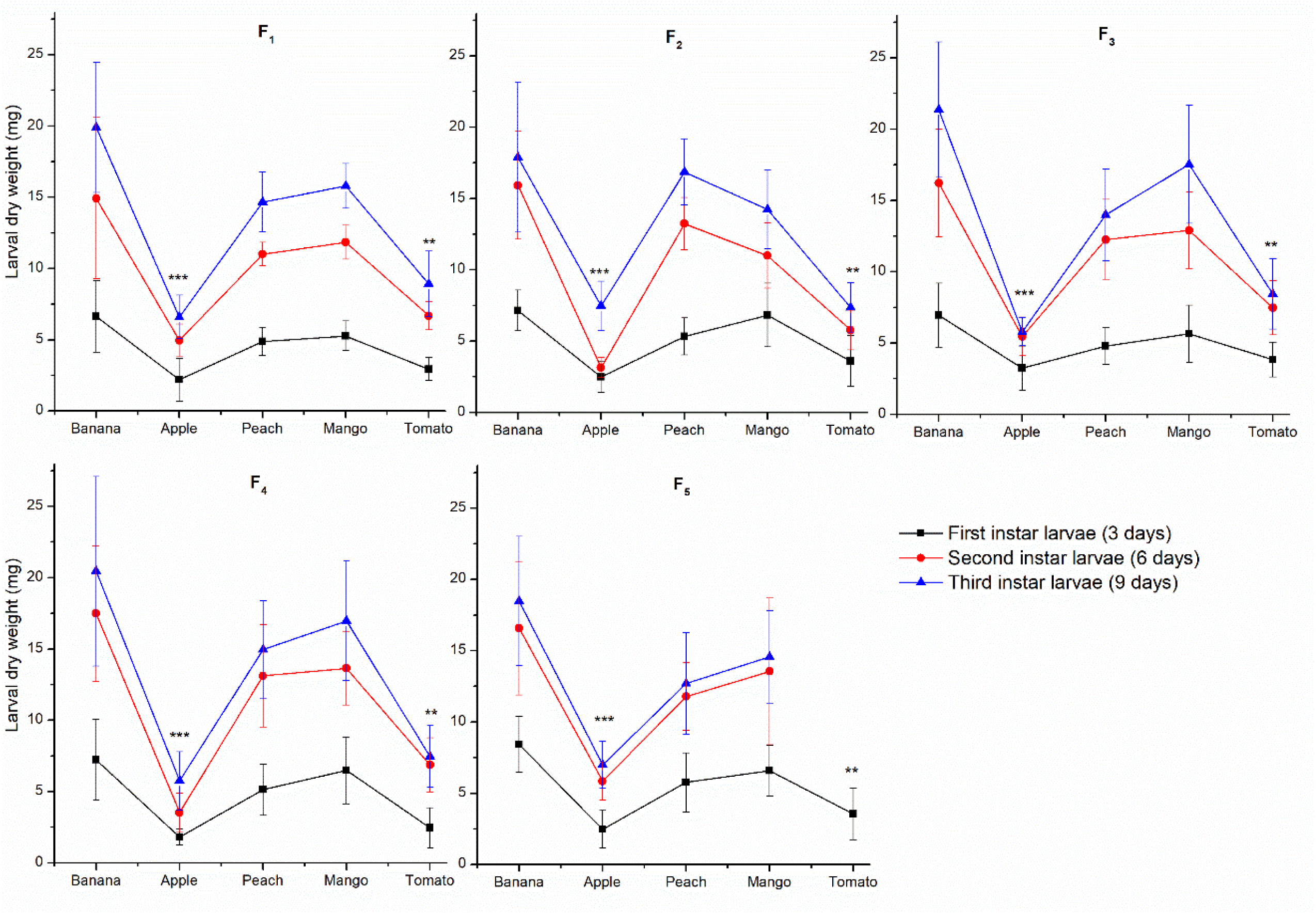
Effect of host fruits suitability on *B. dorsalis* larval dry weight across five generations

#### Developmental growth time of larvae

The larval development (length and weight) was inversely proportional to developmental duration (time from oviposition to adult emergence). The higher the larval length, the shorter the development time and vice versa. The larval development time was significantly extended in apple and tomato (F = 78.384; df = 1, 4; P ˂ 0.001), but was shortened in banana (F = 97.154; df = 4; P ˂ 0.0001) in comparison to peach and mango (Table 1).

**Table 1.** Biological cycle duration of *B. dorsalis* (from first instar larvae to first instar larvae) as affected by host fruit suitability

| Host fruits | Developmental duration (days) |  |  |  |  |
| --- | --- | --- | --- | --- | --- |
|  | F1 | F2 | F3 | F4 | F5 |
| Banana | 28±3<sup>c</sup> | 27±1<sup>c</sup> | 28±2<sup>c</sup> | 27±3<sup>c</sup> | 27±3<sup>c</sup> |
| Apple | 36±4<sup>a</sup> | 36±2<sup>a</sup> | 35±3<sup>a</sup> | 34±5<sup>a</sup> | 35±4<sup>a</sup> |
| Peach | 31±2<sup>b</sup> | 30±2<sup>b</sup> | 31±2<sup>b</sup> | 29±3<sup>b</sup> | 30±2<sup>b</sup> |
| Mango | 30±2<sup>b</sup> | 28±1<sup>c</sup> | 29±2<sup>b</sup> | 31±2<sup>b</sup> | 30±3<sup>b</sup> |
| Tomato | 33±3<sup>a</sup> | 31±2<sup>b</sup> | 32±2<sup>ab</sup> | 30±3<sup>b</sup> | 31±2<sup>b</sup> |

Means followed by different letters in the same column and row are significantly different according to ANOVA and LSD test ( $P < 0.05$ ).

### Gut microbial analyses

#### The diversity of operational taxonomic units (OTUs)

A total of 653,932 raw reads were generated from all samples: 38,467±6564 reads on average for each sample and 30,637 reads based on the minimum number of trimmed sequences from each sample. The OTU composition varied and increased significantly across host fruits and experimental generations compared to the parental fly. Overall, 64 OTUs were detected in the original parent (P), 252 and 427 OTUs in Banana at F_1_ and F_5_ generations (B_1_ & B_5_), respectively, and 421 and 1,059 in Apple at F_1_ and F_5_ generations (A_1_ & A_5_), respectively. The reads length average was 426±4 bp, and the minimum and maximum number of sequence read per OTU were 201 bp and 503 bp, respectively. Although the majority of OTUs from B_5_ were shared among other samples, it contained the highest number of OTUs per sample. Only 27 core OTUs were detected in all samples (Fig 4).

**Figure 4.**
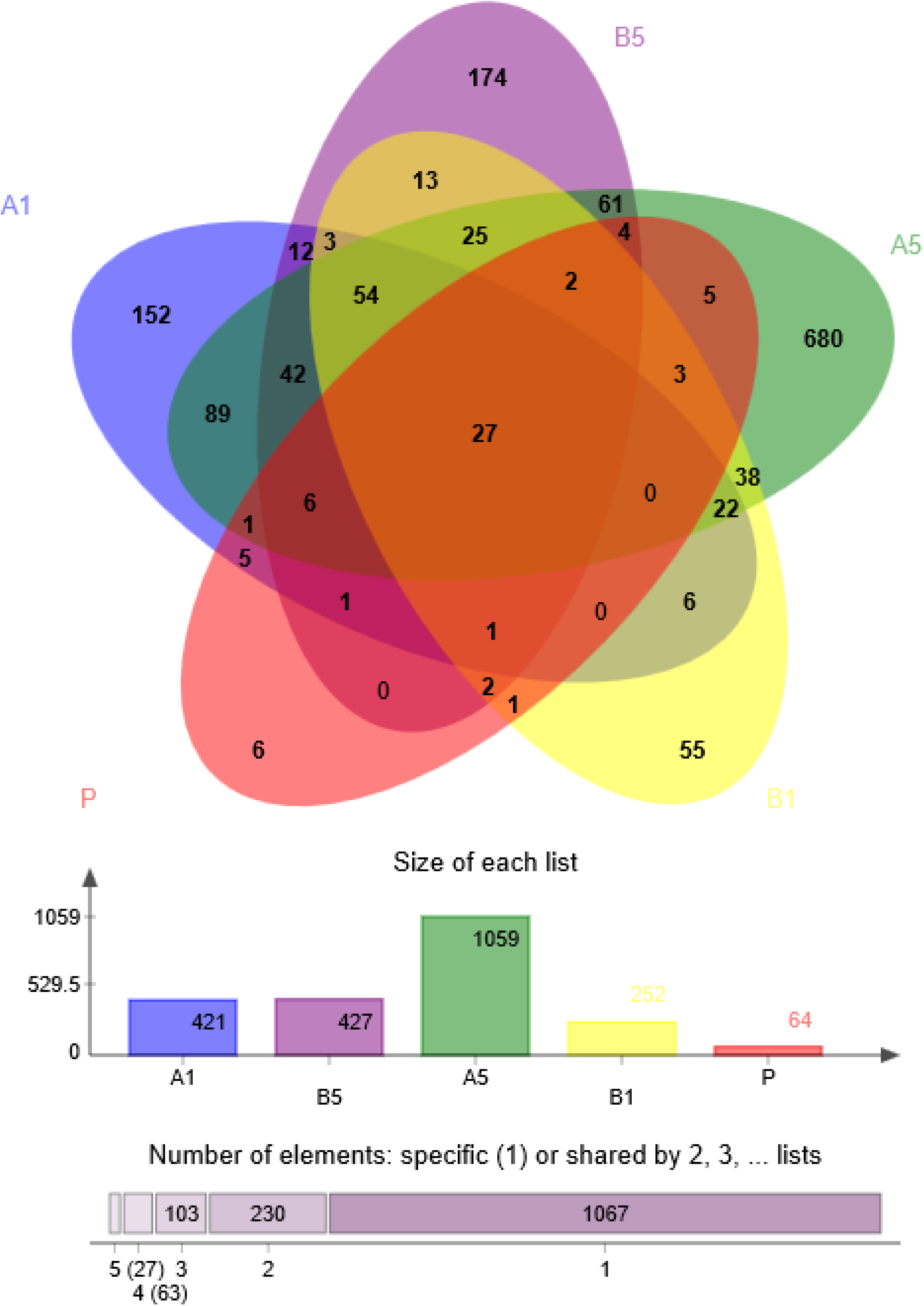
Venn diagram of OTU distribution of sample-specific and shared OTUs. The different colored circles represent different samples or groups. The overlapped areas represent common species among different samples or groups; non-overlapped areas represent unique species in each sample or group.

#### Bacterial richness and diversity

The community diversity (Shannon and Simpson) and richness (sobs, Chao, and ace) evaluated at 97% similarity, showed different comparative trends in the prediction of the number and diversity of OTUs from all samples. The Shannon diversity index provides not only species richness (i.e., the number of species present) but how the abundance of each species is distributed (the evenness of the species) among all the species in the community. Here, we recorded significantly higher bacterial diversity across generations in both tested fruits compared to the parental fly (P) (ANOVA, F = 4, df = 11.173, P = 0.017) (Table 1). For example, the Shannon indices of banana were 0.22±0.26 and 2.5±0.13 at F_1_ and F_5_, respectively (t-test, P = 0.00017±0.0), while those of apple were 1.17±0.23 and 2.5±0.13 at F_1_ and F_5_, respectively (t-test, P = 0.0118±0.02). The comparison between fruits revealed higher bacterial community diversity in apple at F1 (Shannon = 1.17±0.23) compared to banana at the same generation (Shannon = 0.22±0.26) (t-test, P = 0.0095) (Table 1). However, at F_5_, the community diversity was similar in both fruits (t-test, P = 0.3782±0.38).

#### Rarefactions

The rarefaction is a computational analysis of species accumulation based on the repeated re-sampling of all clusters. Based on the good coverage (>0.999) (Table 1), almost all the 16S sequences have been annotated. Moreover, the flattening of the rarefaction curves is an indication that the sampling depth was sufficient enough to detect all OTUs from our samples and an additional sampling effort would only produce few extra OTUs (Fig 5).

**Figure 5.**
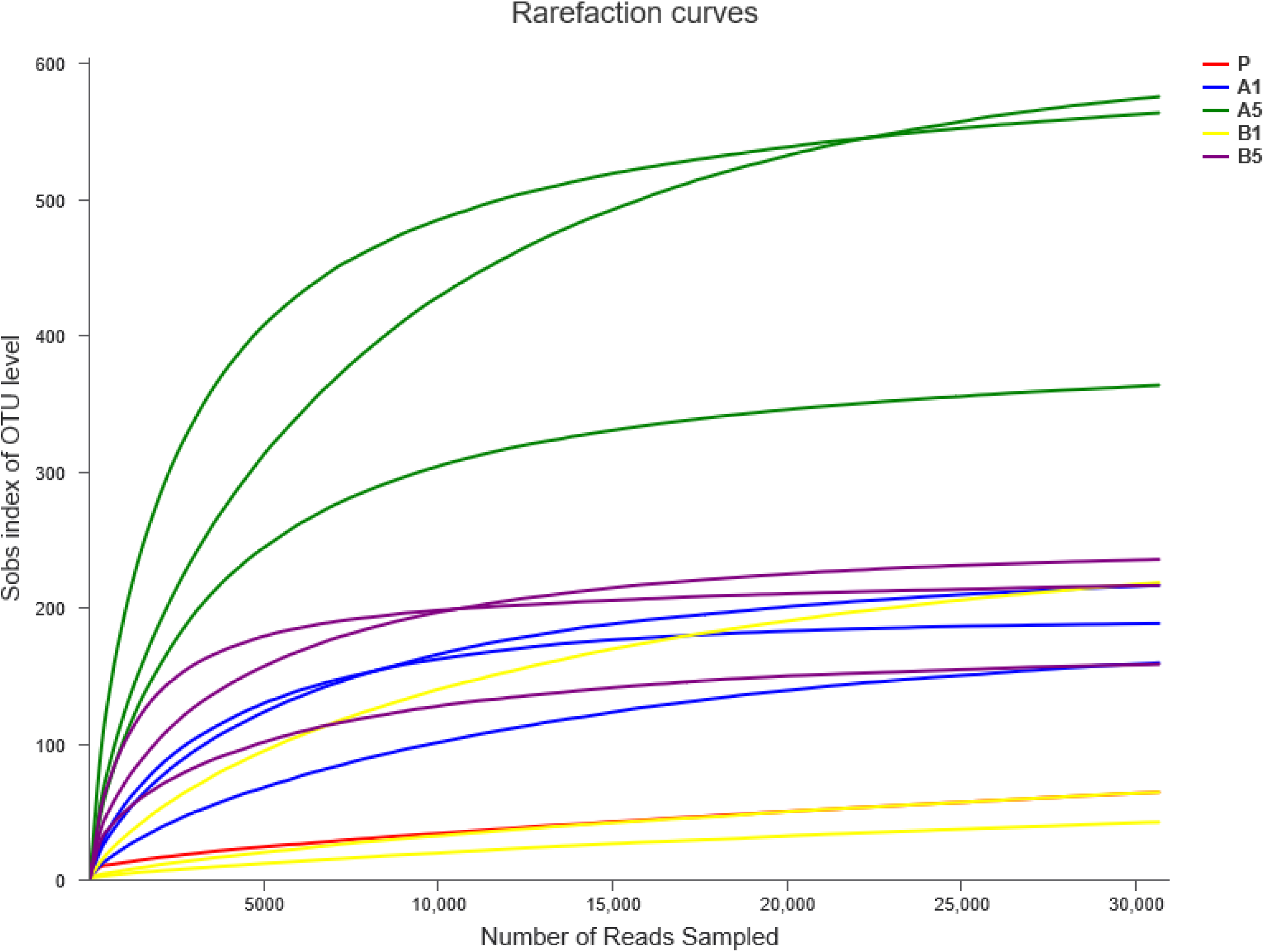
Multiple samples rarefaction curves based on 16S rRNA genes sequencing. The X-axis represents the number of randomly selected sequences; The Y-axis represents the number of species (such as Sobs) or diversity indices (such as Shannon) of each sample. Legends P: parental female; A_1_ & _A5:_ Apple at first and fifth generations; B_1_ & B_5_: Banana at first and fifth generations.

#### Bacterial community composition

The original bacterial community was represented by two phyla only: Proteobacteria accounted for 95.67%, and Firmicutes accounted for 4.19%. The structure of the vast majority of the gut bacterial community did not change at the first generation (F_1_) of both host fruits, but there was an emergence of Bacteroidetes which was not detected from the original community. At F_1_, Proteobacteria represented 96.04% and 99.08% in apple and banana, respectively, and Firmicutes represented 1.64% and <1% in apple and banana, respectively. However, considerable community abundance shift was observed at the fifth generation (F_5_). The Proteobacteria communities dropped to 34.41% and 47.36% in apple and banana, respectively, in comparison with the to F_1_ and P, while the populations of Firmicutes increased considerably at F_5_ (49.82% and 37.11% in apple and banana, respectively). Also, Actinobacteria populations were detected in both fruits (Fig 6). The heatmap showing the relative abundance of taxa in different samples and groups on a certain taxonomic level can be found in Fig S9.

**Figure 6.**
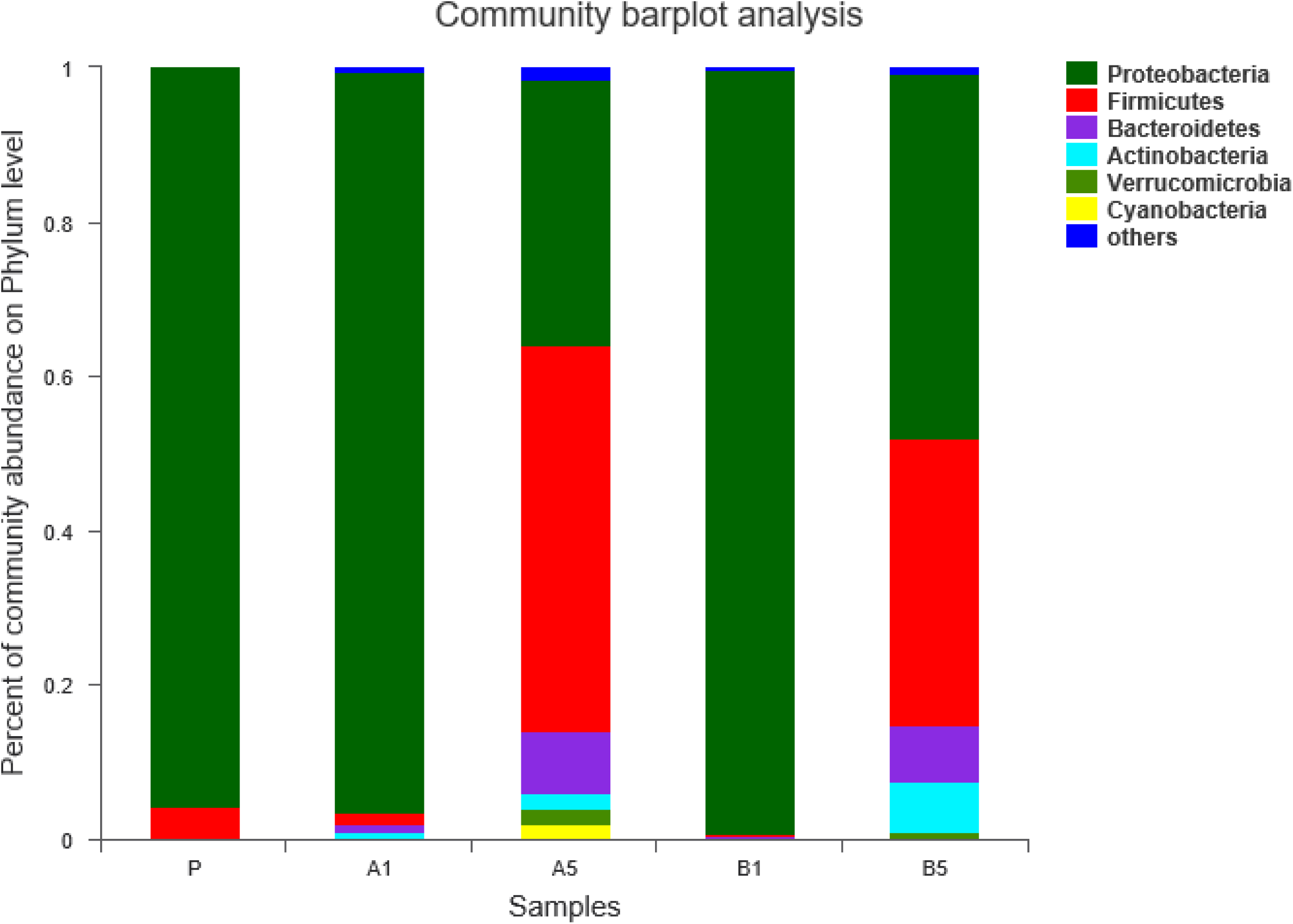
Bacterial community compositions at the phylum level as revealed by high throughput sequencing. Different colored bars represent different species, and the length of the column represents the proportion of the species. Legends: P: parental female; A_1_ & A_5_: Apple at first and fifth generations; B_1_ & B_5_: Banana at first and fifth generations.

Twenty-one bacterial families in total were annotated from all samples, and the number of families increased across generations. Enterobacteriaceae (91.75%) were the most dominant families in the parental female (P), followed by Pseudomonadaceae (3.81%) and Enterococcaceae (2.89%) in lesser proportions. At F_1_, Pseudomonadaceae (67.31%) and Enterobacteriaceae (27.07%) were highly detected in apple while Enterobacteriaceae (97.24%) was predominant in banana (Fig S1, Supplementary). At F_5_, the bacterial families were more diversified in both fruits (12 and 17 in apple and banana, respectively). For instance, Leuconostocacceae (22.66%), Streptococcaceae (19.38%) and Acetobacteriaceae (17.64%) were detected in higher proportions in apple compared to F_1_, while Enterobacteriaceae (27.62%), Bacillaceae (11.93%), Streptococcaceae (10.84%) and Pseudomonadaceae (10.04%) were highly represented in banana compared to their proportions at F_1_ (Fig S1, Supplementary).

At the genus level, *Enterobacter* (85.05%) was dominant in the parental female. At F_1_, *Pseudomonas* (67.31%) and *Enterobacter* (26.68%) were highly represented in apple while *Morganella* (96.82%) was the predominant genus in banana. However, at F_5_, *Lactococcus* (19.12%), *Fructobacillus* (18.58%), *Gluconobacter* (16.99%) and *Morganella* (22.23%) emerged abundant in apple, while *Morganella* (22.23%), *Bacillus* (11.90%), *Lactococcus* (10.74%) and *Pseudomonas* (10.64%) were dominantly represented in banana. Overall, in apple, *Enterobacter* proliferated significantly at F_5_ compared to *Pseudomonas*. However, in banana, Morganella was consistent across generations (Fig S1, Supplementary).

#### Bacterial structure distribution

On the basis of Bray-Curtis distance algorithm, the gut bacterial community structure varied significantly between host fruits and across generations in all samples (Bray-Curtis distance statistics, A = 0.009014; P = 0.0012 & A = 0.000852; P = 0.01) (Fig 7. However, the clustering of different samples into groups indicates that the bacterial communities within groups are highly similar (ANOSIM: R = 0.4259; P = 0.019) (Fig 7).

**Figure 7.**
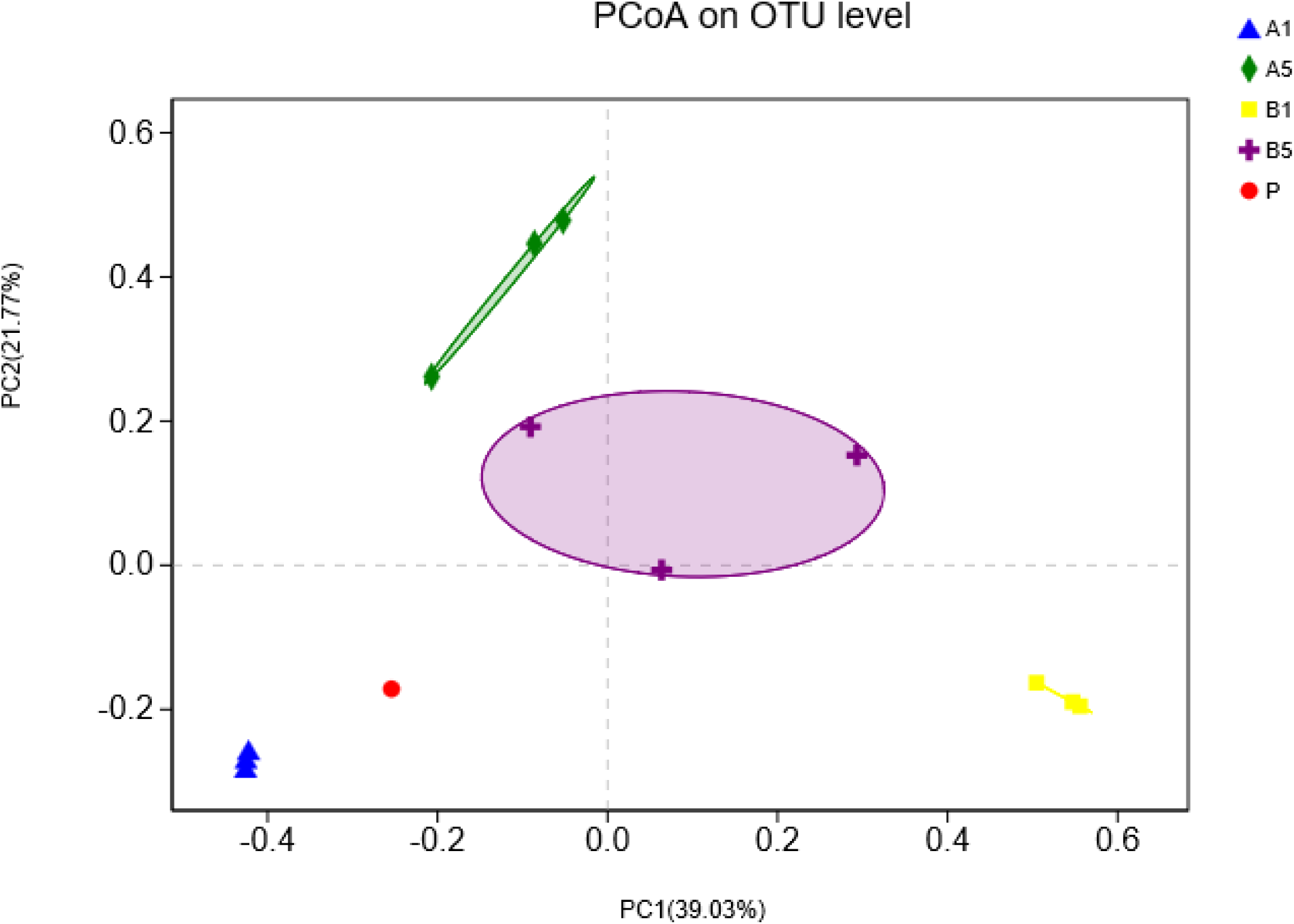
Principal Coordinates Analysis (PCoA) showing the variations of the gut bacterial community of *B. dorsalis* as affected by time and host fruits. Note: The X and Y axes represent two selected principal coordinate components. Legends: P: parental female; A1 & A5: Apple at first and fifth generations; B1 & B5: Banana at first and fifth generations. The percentage “%” indicates the contribution proportion of each principal coordinate component to the sample composition variance. The scales on X-axis and Y-axis are relative distance with no practical significance. Different groups are marked in different colors and shapes. Close samples have a similar community composition structure.

## Discussion

Gut bacteria and fruit flies share close evolutionary relationships which mutually influence the physiology and ecological adaptations of both parties. In this interaction, the larval stages play a dominant role in the survival of the insect whose capacity to stand across multiple generations depends on host suitability for larval growth sustained by its gut microbiome. If fruit fly larvae gain many of their primary nutritional needs from bacterial metabolic activity, then maybe the quality of fruit for fruit fly larvae depends on how good it is for bacterial growth and larval development.

In this study, we assessed how host suitability might shape the larval performance and gut bacterial growth over time. A single female fly was allowed to lay eggs on five different host fruit types and we monitored the larval population dynamics across five generations and bacterial community structure at first and fifth generations.

### Host suitability for larval development

The larval development parameters (length, dry weight, and development duration) increased across larval stages depending on the suitability of host fruits. The highest larval performance was recorded in banana, mango, and peach, while larvae that developed in apple and tomato had the shortest length and dry weight, and extended the development duration across generations. Within the context of adaptive abilities of the fly larvae, these results suggest that the host fruit quality (capacity to offer a readily available source of nutrients) is a critical factor that determines the completion of larval development and its propensity over time [29]. While the adult flies require a diet consisting of protein and sugar for their maintenance, larvae must feed on high sugar content diet for their growth and development [8,10]. Therefore, a suitable substrate for larval growth should be able to offer the necessary carbohydrates to fuel the urgent nutritional requirements of larvae. Larvae under continuous development in tomato did not reach the second instar of their development at the fifth generation. The abortion of larvae may partly be due to the high moisture content in the tomato flesh that led to the drowning of the larvae, and death. In addition, the lack of sufficient sugar metabolites, and free amino acids in tomato may be another cause of the abortion of the larvae over time [30].

### Host suitability for bacterial growth

The gut microbiome plays a prominent role in the nutritional adaptations of insects [18]. When the fly is found in a nutritionally imbalanced environment in terms of nutrients availability, the metabolic activity of gut bacteria may constitute an additional source of nutrients, for their ability to provide amino acids, carbohydrates, vitamins and digestive enzymes to foster and reinforce the adaptive capacity of the host fly [18]. Many previous studies have reported the importance of gut microbiota in host nutrition and physiology [9,10,31-33], and the microbial community divergence at different life stages and geographic areas have been established [8]. However, how the gut bacterial community of larvae developing on different host fruits varies across generations to maintain optimal larval development is not fully understood.

Here, we explored the bacterial community structure and composition of larvae developing in banana (suitable substrate for larval growth and development) and in apple (hostile substrate), highlighted the core bacteria in each fruit and evaluated the extent of their variations across generations compared to the single parental community. Very similar structure of bacterial communities was observed between the parental female and the offspring of the first generation (F_1_), which was dominated by Proteobacteria (≥95%). The Proteobacterial populations decreased significantly, and Firmicutes emerged as the dominant taxa at F_5_. This could be understood at two scales. First, the similarity of bacterial community structure at F_1_ with the parental fly (but different at F_5_) could imply that the community structure changes over time to allow the larvae to adapt in an environment with either no food available or less suitable food for their optimal growth. Secondly, although originated from the same parent, the differences observed between fruits at F_5_ may come at the cost of the different degree of suitability of the substrate and the survival emergency in the long run. Therefore, Proteobacteria and Firmicutes appeared as the driving forces of this behavior due to their prevalence and persistence over time [8,34]. Similarly, the stability of these same bacterial phyla (Proteobacteria and Firmicutes) were shown to be responsible for maintaining optimal larval development across life stages of *B. minax* [29].

Furthermore, when compared to the parental female, huge bacterial community diversity and richness was recorded across fruits (higher in apple at F_1_ but similar in both fruits at F_5_) and across generations (higher at F_5_ in comparison to F_1_). This finding puts to the light the plasticity of gut bacteria which can amend its population to help the host circumventing challenges linked to the fruit phenology and nutrient availability. As shown above, banana and apple offered different larval development facilities. While banana allowed the larvae to develop optimally and grow faster (due to their high sugar content and vitamins), apple extended the larval development time and reduced the measured development parameters. The quality of ingested food influences the structure and composition of gut microbiota in insects, some bacterial taxa are advantaged and increase in size while others are disfavored and maintain their population at the survival threshold (Douglas, 2015). The higher bacterial diversity (with the abundance of the family Pseudomonadaceae and the genera *Pseudomonas*) in apple at F_1_ could be an indication that, at early generation, either the apple offered all the required nutrients for bacterial normal growth, or these gut bacteria possess the ability to digest potential recalcitrant dietary constituents from the fruit. Previously, symbiotic bacteria of the genera *Burkholderia* and *Parabacteroides* were suggested to mediate the degradation of hemicellulose and the recycling nitrogen in *Melolontha hippocastani* [35]. At F_5_, Enterobacteriaceae (genus *Enterobacter*) became the most abundant bacterial community in apple, while the same family Enterobacteriaceae (genus *Morganella*) persisted across generations in banana. This suggests to some extent the dependence of the larvae upon Enterobacteriaceae for their optimal development across generations. In this fashion, the larvae can directly use the available resources from the fruits, or use the byproducts of bacterial metabolic activities to compensate the lack of nutrients from their diet [36]. Therefore, gut bacteria and the substrate suitability are crucial to maintaining a balanced food web and the survival of the host across generations.

Overall, the variability of the gut bacterial community structure and composition in response to different host fruits observed in this study could also be driven by, but not limited to, the changes in the gut transcriptome that might have induced a regulation mechanisms of genes linked to nutrition, growth and detoxification. However, additional experimental steps are needed to provide support for such a conjecture. Such experiments might, for example, include studies where flies developed on one type of fruit for many generations on a particular fruit that gave smaller larvae would continue to show poorer development when transferred to a different fruit. The use of axenic fruit and flies represent another way forward to untangle these interactions. At the very least, our findings add another layer of understanding on the various bacterial compromises that enable *B. dorsalis* to feed in a broad range of host fruits. This may explain (but not limited to) the invasive polyphagous state of the fly.

## Conclusion

In this work, we evaluated how different host fruits can act as a suitable substrate for larval development and gut bacterial growth. By allowing a single female fly to lay eggs on different host fruit, we monitored the bacterial population with regards to the parental ones and determined the patterns of their variations across five generations. The results show that the development of larvae depends on the ability of fruits to offer a suitable substrate or media for gut bacterial growth. As such, banana, mango and peach allowed optimal larval development across generations while apple and tomato were less suitable substrate for the larval growth. Overall, the bacterial community structure and composition was similar at the early stage of the exposure to the fruits (with a predominance of Proteobacteria), but the variations arose over time depending on how suitable the substrate was at availing nutrients for the larvae. This study adds a new stair of understanding of the effects of interactions between gut bacteria and the quality of substrate (fruits) on *B. dorsalis* development.

**Table 2.** Alpha diversity indices, showing the diversity and species richness of gut symbionts of *B. dorsalis* as affected by host fruits and experimental generations.

| Samples | Sobs* | Shannon | Simpson | ACE | Chao1 | Coverage |
| --- | --- | --- | --- | --- | --- | --- |

| **P** | **69<sup>e</sup>** | **1.2236<sup>b</sup>** | **0.4126<sup>b</sup>** | **262** | **133<sup>d</sup>** | **0.99** |
| --- | --- | --- | --- | --- | --- | --- |
| A1 | 188±29<sup>c</sup> | 1.17±0.23<sup>b</sup> | 0.51±0.01<sup>b</sup> | 204±21 | 202±25<sup>c</sup> | 0.99±0 |
| A5 | 500±119<sup>a</sup> | 2.86±0.63<sup>a</sup> | 0.24±0.07<sup>c</sup> | 525±127 | 534±128<sup>a</sup> | 0.99±0 |
| B1 | 108±96<sup>d</sup> | 0.22±0.26<sup>c</sup> | 0.94±0.08<sup>a</sup> | 182±70 | 150±93<sup>d</sup> | 0.99±0 |
| B5 | 203±40<sup>b</sup> | 2.5±0.13<sup>a</sup> | 0.18±0.05<sup>c</sup> | 210±40 | 210±38<sup>b</sup> | 0.99±0 |

P: parental female; A1 & A5: Apple at first and fifth generations; B1 & B5: Banana at first and fifth generations. Different letters within columns are statistically different after Student's t-test at $P < 0.05$ ; (\*): average number of species observed in each sample (alpha diversity).

## Data accessibility

The metagenomic data are pending submission to sequence reads archive (SRA) of GenBank. The project references will be provided as soon as possible.

## Funding

This study was funded by the National Natural Science Foundation of China (31661143045), International Atomic Energy Agency (CRP No. 17153 and No. 18269), Agricultural public welfare industry research supported by Ministry of Agriculture of People’s Republic of China (201503137) and the Fundamental Research Funds for the Central Universities (2662015PY148).

## Competing interests

The authors declare that they have no competing interests.

## Author’s contribution

CYN conceived and designed the study. MA conducted the experiments and wrote the first draft of the manuscript. AM, XR and YW analyzed metagenomics data. XQ and SC helped in statistical analyses. AN edited and revised the manuscript. All authors read and approved this submission.

## Acknowledgements

Authors are grateful to two anonymous reviewers for their insightful comments on the early version of this manuscript.

